# Complementary roles of EPS, T3SS and Expansin for virulence of *Erwinia tracheiphila*, the causative agent of cucurbit wilt

**DOI:** 10.1101/2024.06.24.600446

**Authors:** Jorge Rocha, Lori R. Shapiro, Scott Chimileski, Roberto Kolter

## Abstract

*Erwinia tracheiphila* (Smith) is a recently emerged plant pathogen that causes severe economic losses in cucurbit crops in temperate Eastern North America. *E. tracheiphila* is xylem restricted, and virulence is thought to be related to Exopolysaccharides (EPS) and biofilm formation, which occlude the passage of sap in xylem vessels and causes systemic wilt. However, the role of EPS and biofilm formation, and their contribution to disease in relation to other virulence loci are unknown. Here, we use deletion mutants to explore the roles of EPS, Hrp Type III secretion system (Hrp T3SS) and Expansin in plant colonization and virulence. Then, we quantify the expression of the genes encoding these factors during infection. Our results show that Exopolysaccharides are essential for *E. tracheiphila* survival in host plants, while Hrp T3SS and Expansin are dispensable for survival but needed for systemic wilt symptom development. EPS and Hrp T3SS display contrasting expression patterns in the plant, reflecting their relevance in different stages of the infection. Finally, we show that expression of the *eps* and *hrp*T3SS operons is downregulated in mildly increased temperatures, suggesting a link between expression of these virulence factors and geographic restriction of *E. tracheiphila* to temperate regions. Our work highlights how *E. tracheiphila* virulence is a complex trait where several loci are coordinated during infection. These results further shed light into the relationship between virulence factors and the ecology of this pathosystem, which will be essential for developing sustainable management strategies for this emerging pathogen.

## Introduction

Modern agriculture faces the challenge of increasing crop yields to feed a growing human population (Ray et al. 2013). Demand is met through the engineering of industrial monocultural agroecosystems. Such ecosystems are physiologically and genetically homogeneous, making crop plant populations susceptible to emergence of and invasion by virulent plant pathogens, which utilize diverse virulence factors (Anderson et al. 2004; Stukenbrock and McDonald 2008; Fauquet and Fargette 1990). However, the genetic basis of virulence of agriculturally important non-model plant pathogens is often under-investigated relative to well-studied laboratory models (Mansfield et al. 2012), despite the threat that these plant pathogens pose to food security (Strange and Scott 2005).

*Erwinia tracheiphila* Smith (Enterobacteriaceae), the etiological agent of bacterial wilt of cucurbits, is one agriculturally important plant pathogen where the molecular details of its pathogenesis remain understudied (Shapiro et al. 2016). *E. tracheiphila* infects wild and cultivated Cucurbitaceae host plant species. Early reports suggested that development of wilt symptoms is associated with the accumulation of cells in the xylem, perhaps accompanied by exopolysaccharide (EPS) synthesis and consequent biofilm formation. The biofilm then physically prevents the flow of sap (Yao et al. 1996; Sasu et al. 2010b; Rojas et al. 2013; Shapiro et al. 2014). Three clades in *E. tracheiphila* populations are strongly associated with host specificity towards cucumber, melon, or squash (Shapiro et al. 2018; Rojas et al. 2013); upon infection, susceptible plants develop systemic wilting symptoms that resemble extreme drought stress, and plant death often occurs within 2-3 weeks after symptoms first appear (Shapiro et al. 2012; Sasu et al. 2009, 2010a).

Economic losses related to *E. tracheiphila* are only reported from agroecosystems in temperate Northeastern and Midwestern North America (Rojas et al. 2015). Restriction to temperate regions is mostly attributed to the geographic distribution of its highly specialized vectors, the striped cucumber beetle, *Acalymma vittata* (F.), and the spotted cucumber beetle, *Diabrotica undecimpunctata* howardi (Barber) (Rojas et al. 2015; Fleischer et al. 1999); it could also be due to decreased virulence of *E. tracheiphila* at higher temperatures (Shapiro et al. 2018). Wilting in plants induces emission of volatile compounds that attract vectors; wilting also facilitates feeding from damaged leaves and pathogen acquisition (Shapiro et al. 2012). Then, transmission to healthy plants occurs via the infected frass in feeding wounds (Leach 1964; Mitchell and Hanks 2009) through the leaves or flowers (Sasu et al. 2010b). Notably, disease spread relies on relatively inefficient processes where initial vector colonization requires concentrated feeding and high bacterial dosage, and subsequent plant infection demands repeated contact with multiple infective vectors (Sasu et al. 2010b; Shapiro et al. 2014).

*E. tracheiphila* was first described by the pioneering plant pathologist Erwin Smith in 1911 (Smith 1911). However, molecular tools and protocols for genetic manipulations were only developed in recent years, allowing the identification of virulence factors. In one hand, the Hrp Type III Secretion System (Hrp T3SS) is essential for virulence in all clades, where T3 effectors contribute to host specificity (Shapiro et al. 2018; Nazareno et al. 2016; Olawole et al. 2022, 2021); Hrp T3SS is a key virulence system found in many Gram-negative bacterial plant pathogens (Buttner and He 2009). In addition, a horizontally acquired Expansin-Gh5 protein is essential for bacterial movement in the vascular system and for systemic colonization (Rocha et al. 2020). Expansins are cellulose manipulating, non-enzymatic proteins with contrasting functions in different plant-associated microbes and whose mechanisms for pathogenesis remain poorly defined (Cosgrove 2017; Chase et al. 2020). These facts highlight *E. tracheiphila* as a model with unusual traits in comparison to other vector transmitted and/or xylem restricted bacterial plant pathogens (Olawole et al. 2021). Yet, many questions remain to be addressed in the pathogenesis of *E. tracheiphila,* such as the role biofilm formation in the plant, the interaction and temporal dynamics of different virulence factors during infection, and their respective contribution to colonization and wilt.

In this work, we explore the roles of virulence factors from *E. tracheiphila* BHKY during squash infection, focusing on EPS and Hrp T3SS. First, scanning-electron microscopy confirmed the presence of a bacterial biofilm in the xylem of infected plants. Using deletion mutants, virulence assays, and direct colony forming unit (CFU) counts from plants, we found that EPS is essential for survival in the plant upon inoculation, while Hrp T3SS is dispensable for within-plant survival and growth but required for wilt development, where Hrp T3SS and Expansin act complementarily to effect systemic wilt. Next, inoculations with reporter strains revealed contrasting patterns of expression between EPS and Hrp T3SS during infection. Finally, in vitro assays showed temperature-sensitive expression of *eps* and *hrp*T3SS. Our results shed light on the relationship between virulence factors of *E. tracheiphila* and the unique ecological traits of this non-model bacterium. These results can now help guide the development of sustainable control strategies for bacterial wilt of cucurbits that focus in different stages of disease development and spread.

## Results

### Visualization of *E. tracheiphila* biofilm during squash infection

Wilt development in squash due to *E. tracheiphila* infection has been largely attributed to physical clogging of the plant vascular system related to bacterial growth and biofilm formation that prevent the flow of xylem sap, causing water depravation in vines (Rojas et al. 2015; Leigh and Coplin 1992). Despite these observations, studies are yet to provide direct evidence of biofilm formation during infection. To address biofilm formation during plant colonization by *E. tracheiphila*, leaf petioles of wilting squash seedlings infected with the BHKY wild-type (WT) strain were analyzed at 15 DPI using scanning electron microscopy (SEM). The SEM images (Figure 1) show that the xylem vessels are partially occluded by microcolonies of bacteria, preferentially found attached to the inner wall of the xylem by a thread-like extracellular matrix (ECM). At higher magnification (Fig. 3, center and right panels), the cells and bundles of ECM material were in some cases found stretching across the width of the xylem. These observations support the notion that the production of an ECM and biofilm formation are essential processes during infection by *E. tracheiphila* BHKY.

**Figure 1.**
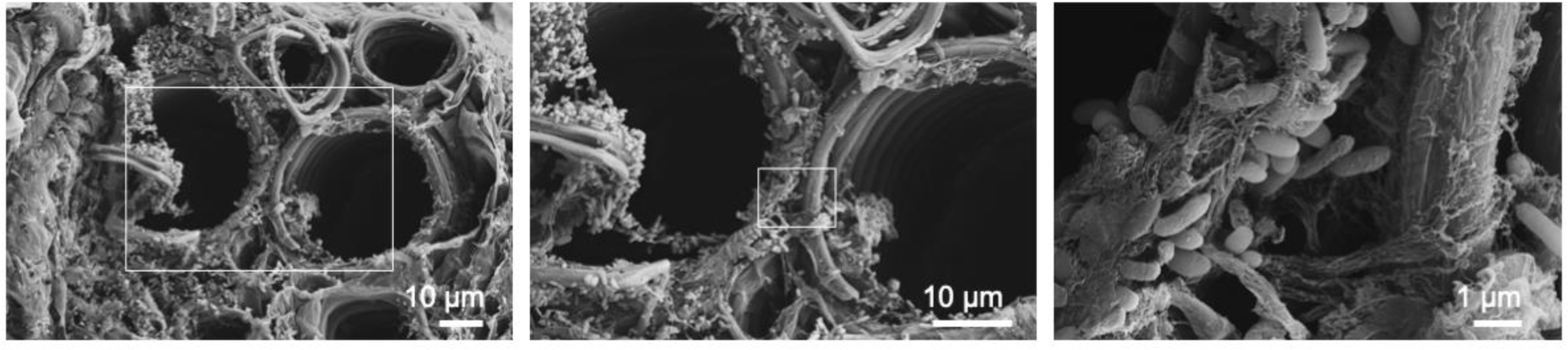
*Erwinia tracheiphila* biofilm-like agglomerations in the vascular system of infected squash plants. Scanning electron microscopy showing the vascular systems of squash plants infected with Et BHKY WT, at 15 DPI. Squares indicate magnification area in the following picture.

### In vitro characterization of Δ*eps* and Δ*hrp*T3SS mutants

Our SEM observations provided direct evidence of a bacterial biofilm in infected squash plants (Figure 1). Interestingly, a locus (*eps*) encoding for the enzymes involved in the synthesis of exopolysaccharide (EPS) was identified in the genome of *E. tracheiphila* strain BHKY (LaSarre et al. 2022; Rojas et al. 2013), with high similarity to virulence loci found in other plant pathogens of the genus *Erwinia* and *Pantoea* (Carlini et al. 2023). In addition, previous work has shown that the Hrp Type III secretion system (Hrp T3SS) and Expansin-GH5 are important virulence factors for *E. tracheiphila* (Rocha et al. 2020; Olawole et al. 2022, 2021).

In order to the study of the participation of EPS during infection, and the complementary roles of the three loci for virulence, we generated clean deletion mutants of the entire *eps* locus, and a large region of the *hrp*T3SS locus in strain BHKY (Fig. 2, Table S1). In vitro, the Δ*eps* mutant displayed reduced colony viscosity, as compared to WT and Δ*hrp*T3SS strains (Supplementary Figure S1). Likewise, a hypersensitivity response assay in tobacco leaves showed that the Δ*hrp*T3SS mutant lost the capacity to induce this response in non-host plants (Supplementary Figure S2), which is directly associated with Type 3 effectors (Nazareno et al. 2016; Cornelis and Van Gijsegem 2000). These assays confirmed that the operons are functional in *E. tracheiphila* BHKY. We described deletion of the *exlx*-*gh5* operon from *E. tracheiphila* BHKY (Figure 2, bottom) and characterization of the Δ*exlx-GH5* mutant in our previous work (Rocha et al. 2020).

**Figure 2.**
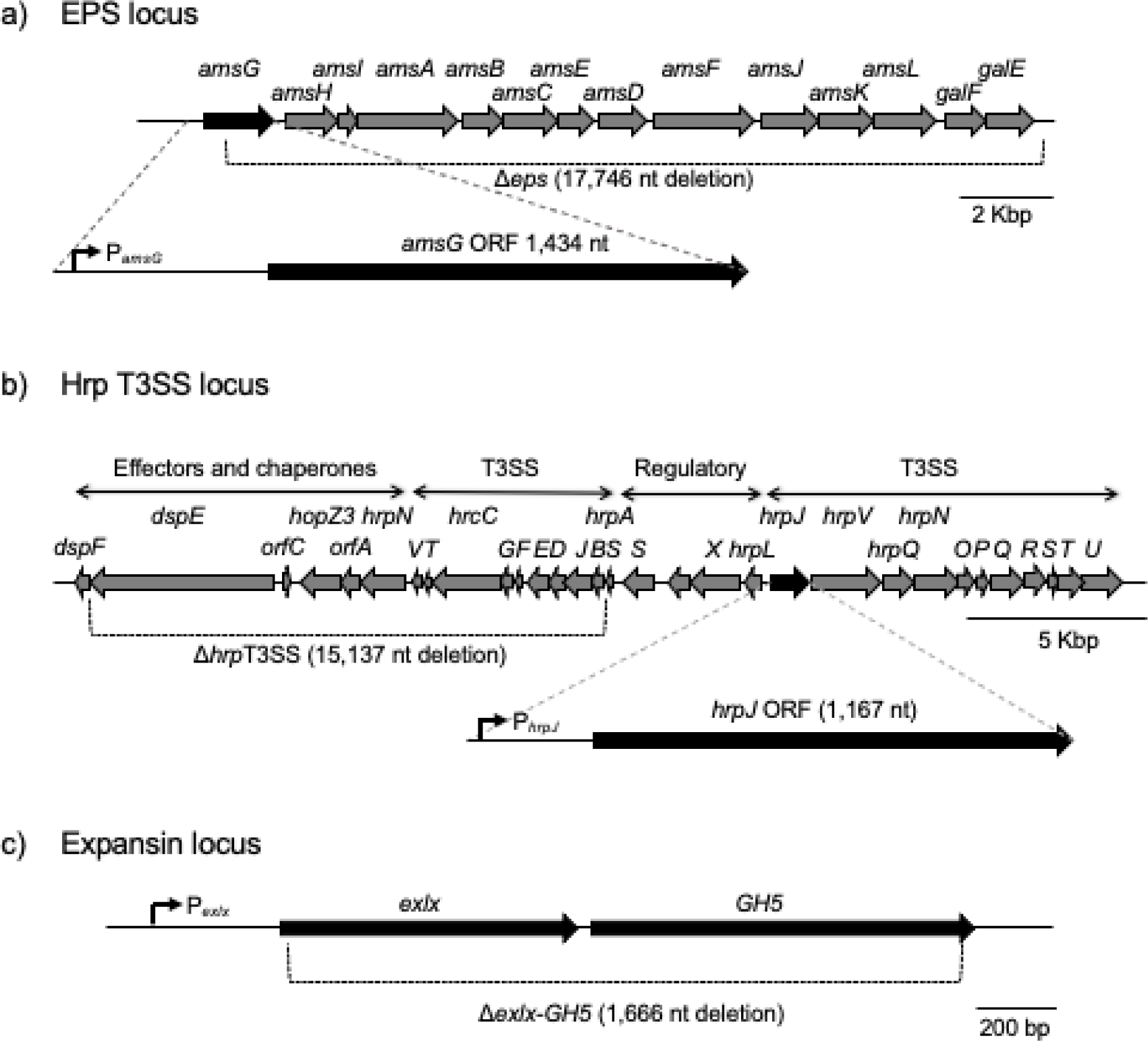
Schematic representation of loci coding for proteins related to (a) Exopolysaccharide (EPS) synthesis, (b) HrpType III Secretion System (Hrp T3SS) machinery, and (c) Expansin. Location of deletion fragments in the corresponding mutants is shown with brackets. Single genes *amsG* and *hrpJ* are shown at higher scale to indicate location of promoter regions (broken arrows) included in the corresponding transcriptional fusions.

EPS is essential for virulence of *E. tracheiphila* strain BHKY.

To determine the contribution of EPS to the virulence of *E. tracheiphila*, squash plants (*Cucurbita pepo* ‘Yellow Crookneck’) were inoculated with the WT or the Δ*eps* mutant strain. All plants inoculated with the WT strain developed symptoms by 12 days post inoculation (DPI). By 21 DPI, 9 out of 12 plants inoculated with WT had died. No wilting symptoms or death were observed for plants inoculated with the mutant strain. Thus, the Δ*eps* strain is avirulent (Figure 3).

**Figure 3.**
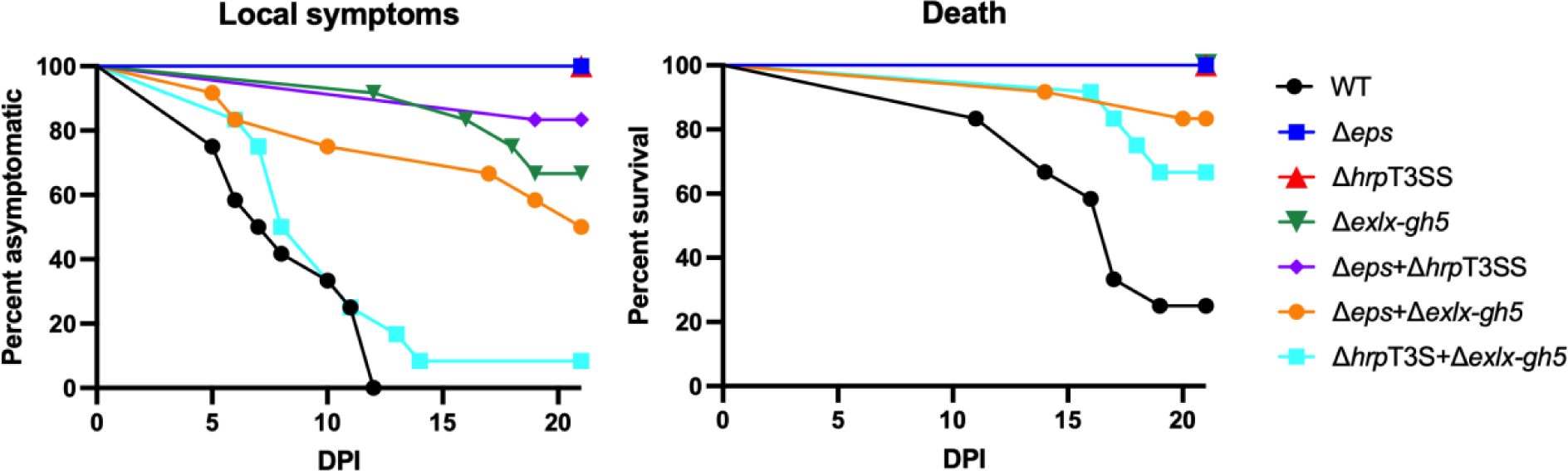
Virulence of *E. tracheiphila* BHKY WT and deletion mutants in single- and co-inoculations. Plants (n=12) were inoculated with WT, mutants or pairwise combination of mutants and virulence was followed during 21 DPI by assessing detection of local symptoms (left) and plant death (right).

Next, we performed pairwise co-inoculations of squash plants with Δ*eps*, Δ*hrpT3SS* and Δ*exlx-gh5*, in order to explore if there was any extracellular complementation between pairs of mutant strains, which would be indicative of an extracellular function. For this, we tested the virulence of Δ*hrp*T3SS and Δ*exlx*-*GH5* mutant strains. Previous work showed that Hrp T3SS is essential for virulence of *E. tracheiphila* (Olawole et al. 2021), while Expansin-GH5 is important for systemic wilt (Rocha et al. 2020); we confirmed those phenotypes in single strain inoculations (Figure 3). In the case of mixed inoculations, the Δ*hrpT3SS* and Δ*exlx-gh5* pair resulted in increased symptoms and death of plants, compared to the individual inoculations (Figure 3, Supplementary Table S2), indicating that both virulence factors act extracellularly on plants since *hrp*T3SS is provided by the Δ*exlx-gh5* mutant and *vice versa*. Co-inoculations of the Δ*eps* mutant with either Δ*hrp*T3SS or Δ*exlx-gh5* did not result in extracellular complementation, indicating that EPS produced by Δ*hrp*T3SS or Δ*exlx-gh5* cannot be used by the Δ*eps* mutant. Overall, our results show that *E. tracheiphila* pathogenesis in squash plants is a complex trait involving multiple loci, where Hrp T3SS and Expansin act complementarily on the host plants

Colonization dynamics of *E. tracheiphila* Δ*eps* and Δ*hrp*T3SS mutants during squash infection.

Our previous work showed that Expansin-GH5 is important for systemic colonization, and in turn, for systemic wilt of squash plants (Rocha et al. 2020). To test whether the avirulent phenotypes of Δ*eps* and Δ*hrp*T3SS mutants are related to a growth defect of these strains in the plant, we followed colonization after squash inoculation. Plants were inoculated with 10^7^ cells of individual strains, and CFUs were measured in the site of inoculation at 2, 6, 12 and 18 DPI. The WT strain was found at ≍2×10^7^ cells per g of fresh plant at 2 and 6 DPI, and reached a maximum colonization of ≍ 3×10^9^ CFU/g at 12 DPI (Figure 4).

**Figure 4.**
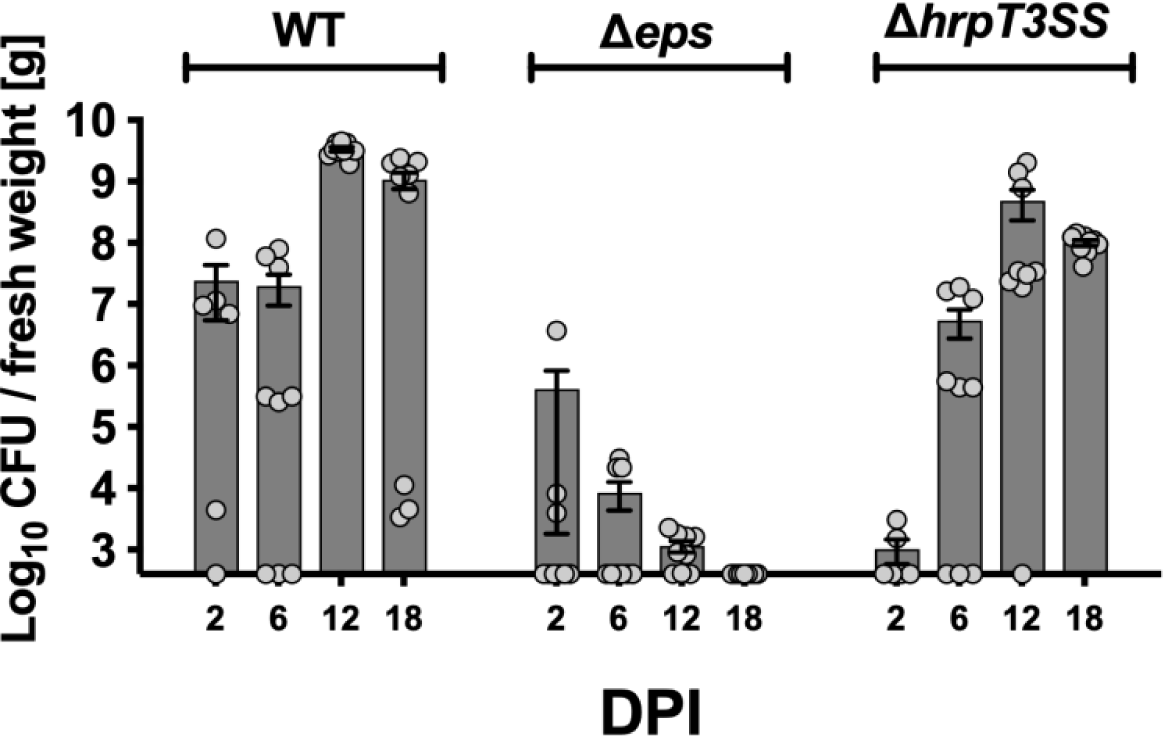
Colonization of *E. trachiephula* WT, Δ*eps* and Δ*hrp*T3SS strains in squash plants. Plants were inoculated in the petiole of the second true leaf, and CFU counts were carried out in the site of inoculation. Bars represent average ± Standard Error of the Mean and gray dots indicate individual biological replicates (n=6-9)

Growth of *E. tracheiphila* in the plant was severely affected by the deletion of *eps* operon. The Δ*eps* mutant was detected from inoculated plants at ≍ 4×10^5^ CFU/g at 2 DPI, and decreased in later time points until it was undetectable in most plants at 12 and 18 DPI. In contrast, Δ*hrp*T3SS mutant strain was able to grow in the plant at similar levels than the WT strain, reaching a maximum of ≍ 4×10^8^ CFU/g at 12 DPI (Figure 4). Noteworthy, robust growth of the Δ*hrp*T3SS mutant in the site of inoculation occurs in the absence of wilting symptoms (Figure 2), as opposed to the Expansin deletion mutant Δ*exlx-gh5* where growth and wilting symptoms are detected in the inoculation site (Rocha et al. 2020). In vitro, both Δ*eps* and Δ*hrp*T3SS grew robustly at only a slightly slower rate compared to the WT strain (Supplementary Figure S3). In summary, these results show that EPS is essential for survival in the plant, while Hrp T3SS is dispensable for growth but necessary for wilt symptoms, and Expansin mediates movement in the plant for systemic infection.

### Validation of fluorescent reporter strains

We studied the expression of virulence loci using transcriptional fusions with *gfp* (see methods and Figure 1). For the EPS reporter, the *gfp* gene is controlled by the P*_amsG_* promoter; for the T3SS reporter, *gfp* is directed by P*_hrpJ_*; for Expansin, we used the P*_exlx_* (Figure 1, Supplementary Table S1). In vitro, *amsG* was highly expressed in rich KB media (Figure 5a), while the *hrpJ* promoter is not active in these conditions (Figure 5b). Expression of *hrpJ* expression was detected only in Induction Media (IM) which simulates the conditions found in the xylem (Wei et al. 1992), with a maximum peak at 10 h of incubation (Figure 5c). Fluorescence in vitro was not detected from the Expansin reporter strain with the P*_exlx_’gfp* transcriptional fusion in either rich KB media or IM (Supplementary Figure S4).

**Figure 5.**
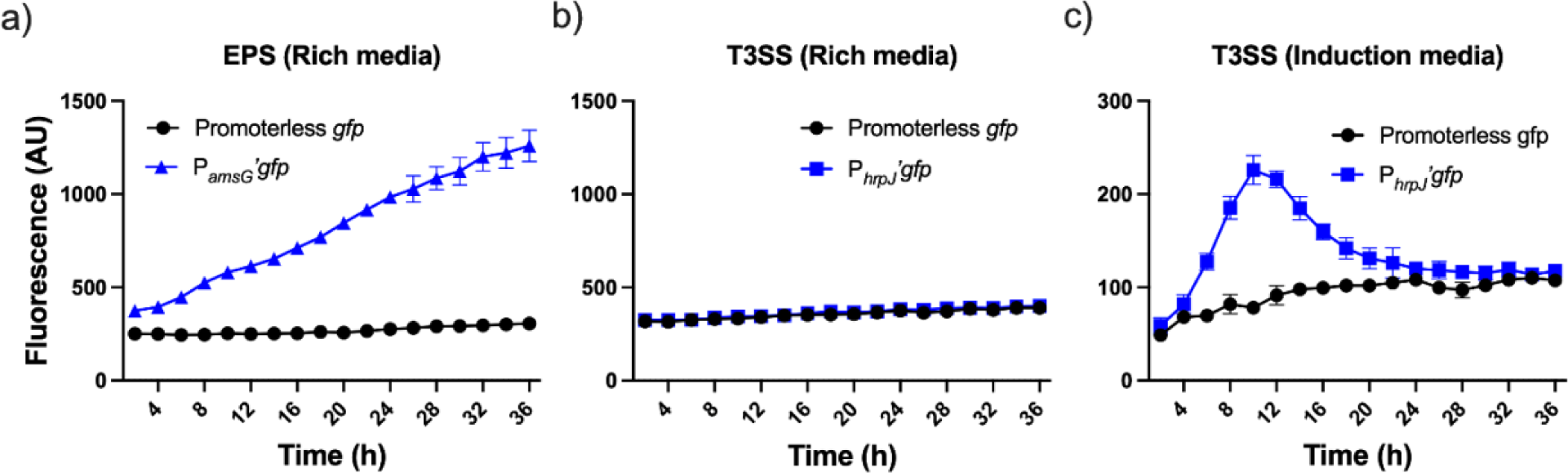
Expression of *amsG* and *hrpJ* in liquid cultures. *E. tracheiphila* reporter strains for EPS (P*_amsG_*’*gfp*) and hrpT3SS (P*_hrpJ_*’*gfp*) or promoterless gfp control were grown in rich media or induction media, and GFP fluorescence was followed for 36 h.

Additional to GFP, these reporter strains carry a constitutively expressed mCherry gene, which allow parallel visualization of colonization and expression of virulence factors directly from infected plants. Squash plants were inoculated with P*_amsG_*’*gfp* or P*_hrpJ_*’*gfp* reporter strains or the promoterless *gfp* control strain, and petiole samples from the site of inoculation of symptomatic plants were analyzed at 15 DPI using fluorescence microscopy. *E. tracheiphila* cells were detected through mCherry fluorescence in the control and reporter strains, mainly in the xylem vessels (Figure 6, top panels). GFP fluorescence was detected from the reporter strains P*_amsG_*’*gfp* and P*_hrpJ_*’*gfp,* indicating that cells are actively synthesizing components related to synthesis of EPS and T3SS machinery at 15 DPI in symptomatic plants (Figure 6). Notably, merged images revealed that GFP signals from *amsG* and *hrpJ* promoters were only partially overlapped with constitutive mCherry signals, indicating that expression of virulence genes is only carried out by a subpopulation. Expression from the *exlx* promoter was not detected in fluorescence microscopy analyses (not shown).

**Figure 6.**
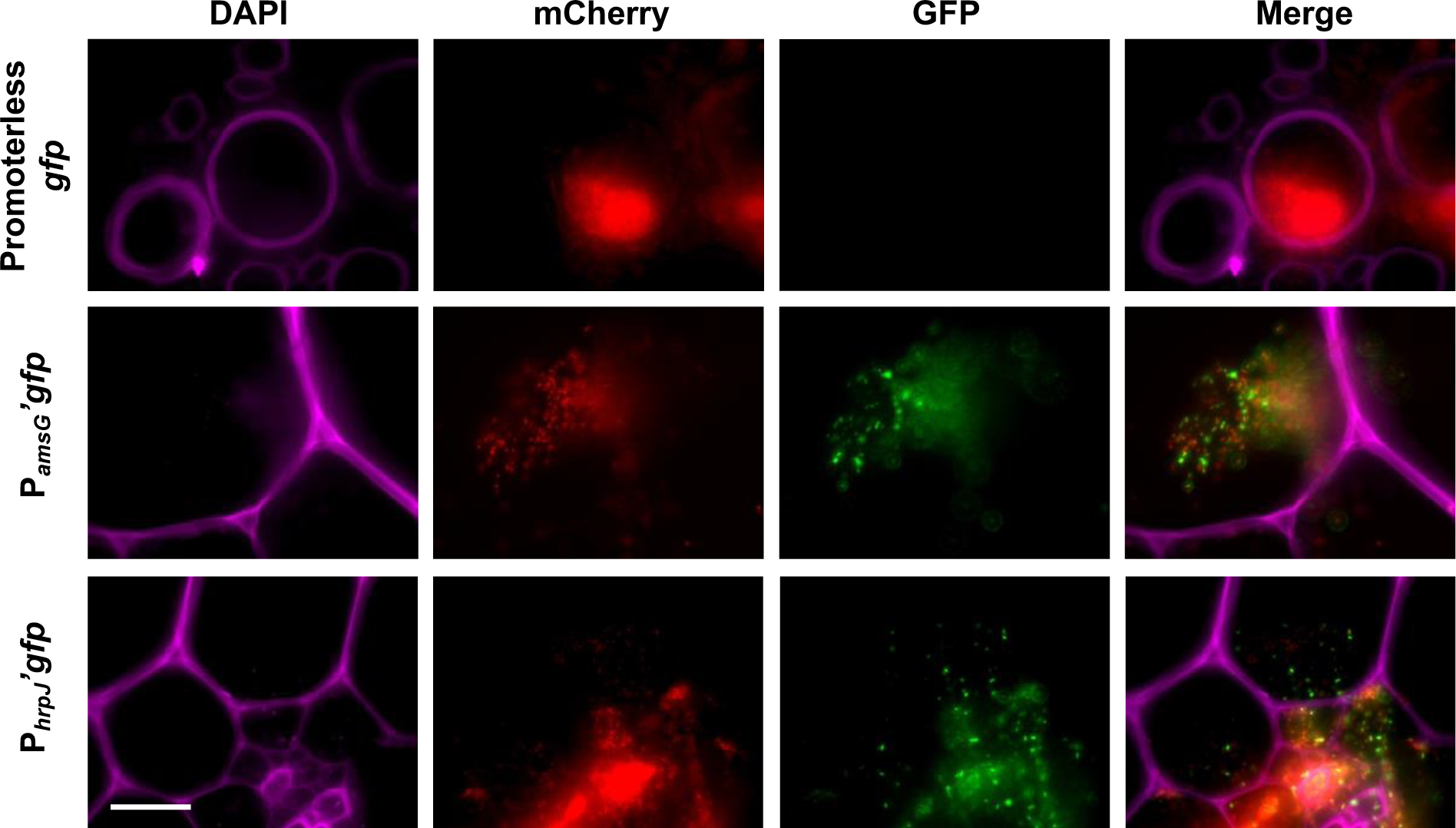
EPS and hrpT3SS are expressed during the infection of *E. tracheiphila* in squash plants. Fluorescence microscopy images were obtained from samples of plants infected with *E. tracheiphila* reporter strains P*_amsG_*’*gfp*, P*_hrpJ_*’*gfp*, and the negative control with promoterless *gfp*. Separate signals are shown for DAPI (plant vascular system), mCherry (constitutive expression in reporter strains), and *gfp* (specific gene expression), as well as the merged image. Pictures were taking at 63× magnification; Scale bar, 50 µm

### Expression patterns of *amsG* and *hrpJ* during infection

Our fluorescence microscopy analyses suggested that expression of virulence factors could be regulated by controlling the percentage of expressing cells during infection. To determine quantitatively the temporal patterns of expression of EPS and Hrp T3SS at the single-cell level, we used flow cytometry. Bacterial cell suspensions were obtained from plants inoculated with the reporter strains at 2, 6, 12 and 18 DPI, and percentage of GFP-expressing cells were calculated from the total population of mCherry-expressing cells.

The gene *amsG* from the EPS operon is expressed in >90% of cells in the inoculum culture (Figure 6a) and this high percentage of expressing cells was maintained at 2, 6, and 12 DPI. In later stages of the infection, expression of *amsG* decreased to <10% of cells. The gene *hrpJ* from the hrpT3SS operon is expressed in <1% of cells in the inoculum but it is rapidly induced in >70 % of cells at 2 DPI (Figure 6b); then, expression decreases to ≍50% of cells at 6 DPI, and ≍15% at 12 and 18 DPI, respectively. We investigated the rapid induction of *hrpJ* gene expression by inoculating plants with the P*_hrpJ_*’*gfp* reporter and the promoterless *gfp* control strains, obtaining cell suspensions after 20 min, 2 h, 6 h, 12 h and 24 h. An increase in percentage of GFP-expressing cells in the reporter strain was first detected at 6 h after inoculation (Figure 6c), suggesting that *E. tracheiphila* responds to the plant host by quickly inducing expression of genes for assembling the *hrp*T3SS machinery. Fluorescent cells were not detected in suspensions obtained from plants inoculated with the Expansin reporter strain with the P*_exlx_*’*gfp* transcriptional fusion (Supplementary Figure S4); Expansin expression may be specialized to a very small subpopulation and/or specific plant locations, or it requires very low protein amounts for functioning.

**Figure 7.**
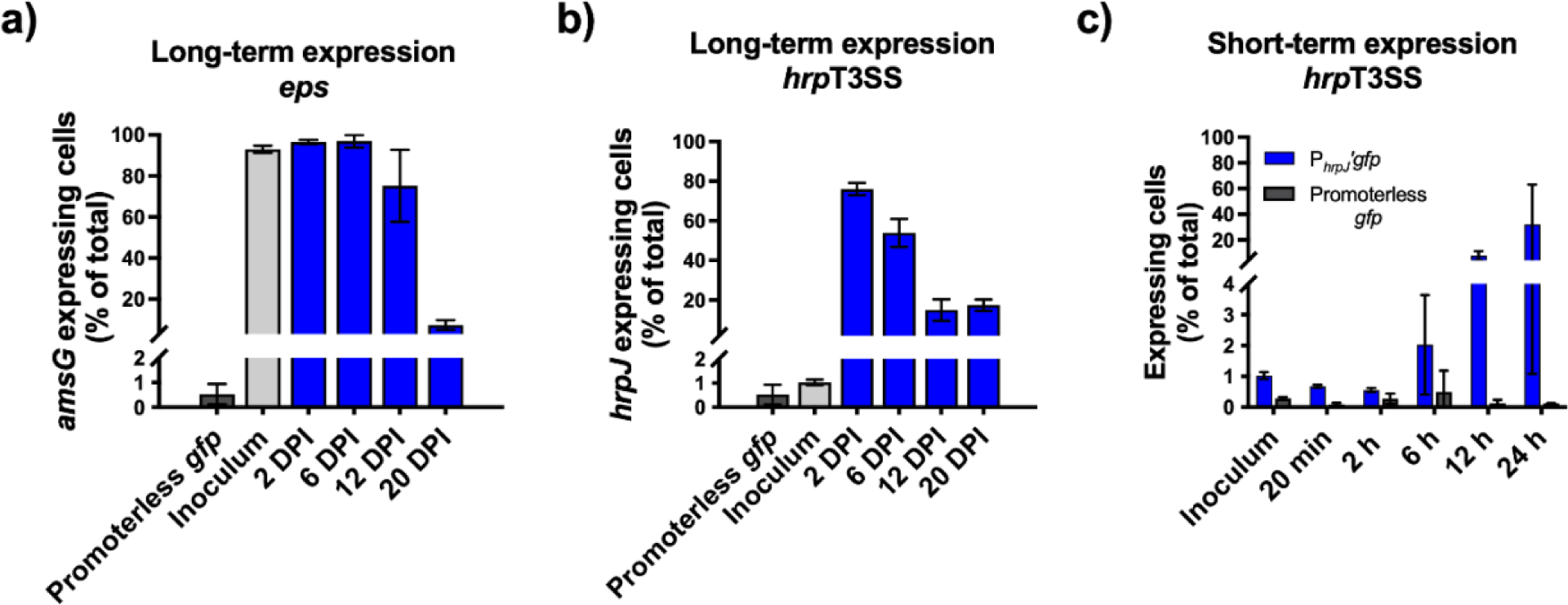
Temporal expression patterns of virulence genes associated to EPS and hrpT3SS from *E. tracheiphila* during squash infection. Expressing cells (%) were quantified by flow cytometry in suspensions obtained from samples of plants infected with *eps* (a), or *hrp*T3SS (b, c) reporter strains (WT genetic background). For reference, expression was analyzed in the overnight liquid culture of the reporter strains and the promoterless gfp control in KB media upon inoculation.

### Thermosensitive expression of *amsG* and *hrpJ*

The limited biogeography of cucurbit wilt caused by *E. tracheiphila* to temperate areas of Northwest and Midwest US has been associated to the distribution of the obligate insect vectors (Rojas et al. 2015; Fleischer et al. 1999). However, plant inoculation experiments in controlled laboratory conditions showed that higher temperatures are related to decreased virulence of *E. tracheiphila* (Shapiro et al. 2018). In order to further understand the effect of temperature on the virulence of *E. tracheiphila*, we investigated expression of *amsG* from the *eps* operon and *hrpJ* from the *hrp*T3SS operon in standing liquid cultures in 25 °C or 30°C. Our results show that the higher temperature decreased the expression of both *amsG* (Figure 8a) and *hrpJ* (Figure 8b). The decreased expression is not a result of reduced growth, since *E. tracheiphila* BHKY WT strain grew similarly in both temperatures (Supplementary Figure S5). These results indicate that thermosensitive expression of EPS and hrpT3SS in *E. tracheiphila* could contribute to geographic restriction of cucurbit wilt to temperate regions.

**Figure 8.**
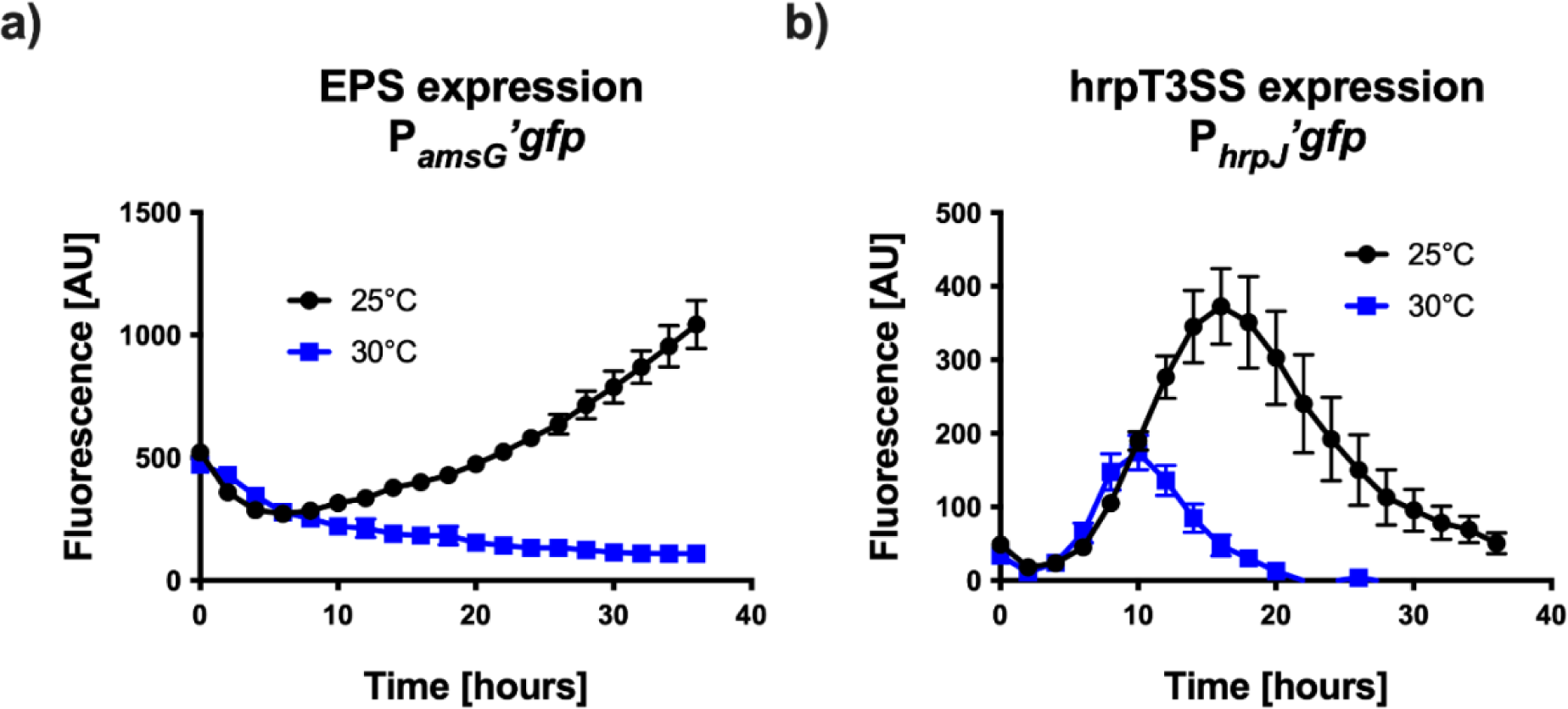
In vitro expression of genes related to a) EPS and b) hrpT3SS is sensitive to increased temperature. *E. tracheiphila* reporter strains P*_amsG_*’*gfp* and P*_hrpJ_*’*gfp* were grown in KB rich media or Induction Media, respectively; GFP fluorescence was followed for 36 h at 25°C and 30°C. Average ± standard deviation of six replicates is shown.

## Discussion

We investigated the complementary roles of *eps*, *hrp*T3SS, and *exlx-GH5* loci from *E. tracheiphila* during squash infection by exploring their relevance for plant colonization and wilt induction, expression patterns during infection, and effect of temperature for expression in vitro. Previous studies showed that hrpT3SS and Expansin are important virulence factors of *E. tracheiphila* (Rocha et al. 2020; Olawole et al. 2021), challenging the previous notion that wilt was caused solely by bacterial accumulation, xylem clogging, and prevention of sap flow in the vascular system (Yao et al. 1996; Sasu et al. 2010b; Rojas et al. 2013; Shapiro et al. 2014). Here we show that at least three virulence determinants contribute with contrasting roles and expression patterns that are required for survival in the plant and biofilm formation (EPS), systemic colonization (Expansin), and wilt (Hrp T3SS).

Biofilm formation is a very important trait for plant pathogens, mediating xylem and vector colonization (Danhorn and Fuqua 2007; Mina et al. 2019). Exopolysaccharide (EPS) is a major biofilm component that can be found as an attached cell capsule or as part of a shared the extracellular matrix (ECM) (Denny 1995; Whitfield et al. 2020). Our results show that the *eps* locus is essential for survival of *E. tracheiphila* in the plant after inoculation. In early stages of infection, all infecting cells are actively expressing the *amsG* gene since EPS cannot be obtained from neighboring cells, as shown in our co-inoculation experiments. In later stages, only a fraction of bacterial cells expresses *amsG,* and an ECM was directly observed in wilting plants. These results indicate that EPS is initially important for self-protection of each cell from plant defenses, and later in the infection it accumulates in the xylem as part of the ECM that holds the biofilm. The *eps* locus of *E. tracheiphila* is orthologous to the loci related to the synthesis of EPS by the taxonomically close plant pathogens *Erwinia amylovora* and *Pantoea sterwarti* (Carlini et al. 2023). In these model species, EPSs (amylovoran and stewartan, respectively) carry out similar functions impacting virulence, biofilm formation, systemic colonization, and protection from plant defenses (Piqué et al. 2015; Oh and Beer 2005; Bellemann and Geider 1992; Menggad and Laurent 1998; Koczan et al. 2011; Ordax et al. 2010; Dolph et al. 1988; Koutsoudis et al. 2006; Roper 2011). However, the *eps* locus from *E. tracheiphila* has a unique genetic organization, probably related to adaptation in its unique ecosystem (Carlini et al. 2023). In this sense, EPS from *E. tracheiphila* could also contribute to functions relevant for xylem restricted or vector transmitted plant pathogens, such as formation of a structured biofilm (Rapicavoli et al. 2018), clogging of the vascular system (Roper et al. 2007), or vector colonization and transmission to healthy plants (Killiny et al. 2013).

The recent development of molecular tools in *E. tracheiphila* has allowed for the generation of laboratory models to understand the molecular basis of virulence (Vrisman et al. 2016, 2018; Olawole et al. 2021; Rocha et al. 2020). However, virulence is the result of coordinated functions that enable a pathogen to cause harm in its host, which involve processes related to host infection, but also to vector colonization, transmission, or adaptation to environmental conditions (Pedroncelli and Puopolo 2023; Mattock and Blocker 2017; Bucci 2018). By analyzing the effect of gene deletion on virulence and colonization, as well as expression dynamics in vitro and during colonization, here we disentangle the complementary roles of three virulence loci. EPS was essential for survival with high expression levels in rich media and during most of the infection. Constitutive expression of P*_amsG_* in rich media suggests that the EPS is also relevant for bacterial fitness outside of the plant host, (e.g., for host colonization or field transmission) (Killiny et al. 2013; Rapicavoli et al. 2018; Shapiro et al. 2014). Type 3 effectors have been shown to mediate host specificity in *E. tracheiphila* (Shapiro et al. 2018; Olawole et al. 2021). Expression of P*_hrpJ_* (from the *hrp*T3SS locus) was repressed in rich media and induced in the plant 6 h after inoculation. This locus was dispensable for growth in xylem, but essential for wilt development. Notably, a second *inv/spa* T3SS is present in the genome of *E. tracheiphila* (Shapiro et al. 2016), which could contribute to vector colonization (Correa et al. 2012). Finally, Expansin-GH5 protein, which is necessary for systemic colonization of the host plant (Rocha et al. 2020), acts synergistically with Hrp T3SS for systemic wilt, which is in turn important for vector attraction, for facilitating the feeding of vectors from plants, and for effective transmission to healthy plants (Rocha et al. 2020; Shapiro et al. 2014, 2012). Together with previous findings, our results offer an integrated view of the relative contribution of virulence loci for the biology and ecology of *E. tracheiphila.* These findings could guide the development of sustainable and effective control strategies (Pedroncelli and Puopolo 2023; Vrisman et al. 2020) that are directed against specific virulence determinants (Yang et al. 2014; Hotinger et al. 2021; Charro and Mota 2015) for efficiently targeting different stages of plant infection, field transmission, or the lifecycle of the vector.

Pathogen emergence refers to adaptations that result in increased virulence, host range, or geographic scope of a virulent species (Cleaveland et al. 2007; Polgreen and Polgreen 2017). Practices of modern agriculture are highly conductive to the emergence of plant pathogens, which represent a major challenge for food production that relies on intensive systems (Sacristán et al. 2021). For this reason, understanding the molecular, ecological, and evolutionary basis of pathogen emergence in agroecosystems is central for its prevention (McCann 2020). *E. tracheiphila* is an example of pathogen emergence due to human activities, since the introduction of cucumber to America was likely the main factor driving its adaptation from native plants to a new host and subsequent diversification into pathovars (Shapiro et al. 2018). Additionally, emergence into new geographic areas should be a major concern. The disease caused by *E. tracheiphila* is highly restricted to temperate regions of North America due to the geographic distribution of its insect vectors (Rojas et al. 2015; Fleischer et al. 1999). However, our results show that thermosensitive expression of the essential virulence factors EPS and hrpT3SS may be directly related to reduced virulence at higher temperatures (Shapiro et al. 2018) and in turn to geographic restriction of the disease. It is easy to speculate that other human-driven phenomena – such as climate change – could modify the geographic incidence of *E. tracheiphila* in the short term, making it an important risk in non-affected areas (Raza and Bebber 2022; Bebber 2015; Velásquez et al. 2018). Moreover, the genome of *E. tracheiphila* is highly prone to mutations, pseudogenizations and horizontal gene acquisitions (HGT) (Shapiro et al. 2016). Indeed, acquisition of virulence factors through HGT such as cluster-specific T3 effectors (Shapiro et al. 2018, 2016), or the unique expansin-GH5 operon architecture (Rocha et al. 2020; Chase et al. 2020), could be related to the recent emergence of *E. tracheiphila* as a pathogen. Similar events could mediate modifications in the genetic regulatory circuits that result in increased virulence at higher temperature and emergence of this disease in new areas.

## Conclusion

The development of genetic tools to investigate the molecular basis of plant-pathogen interactions for non-model, agriculturally relevant microbes is essential for achieving efficient interventions in the ecology of a pathosystem (Sundin et al. 2016). Although *E. tracheiphila* was one of the first plant pathogens identified, only in recent years, the generation of genetically modified strains and other molecular tools has allowed the study of the molecular basis of virulence. Together with recent studies and earlier field observations, our work shows that virulence of *E. tracheiphila* is the result of a complex interplay between complementary virulence factors such as Hrp T3SS, Expansin, EPS, and biofilm formation, which impact plant colonization and development of disease symptoms. Moreover, these and other virulence determinants impact host range, biogeography, transmission to and from the insect vectors, and disease spread. We expect that understanding of the genetic basis for virulence of *E. tracheiphila* will aid in the search for new molecular targets for biocontrol and ultimately the development of sustainable control strategies that are currently absent. Finally, the *E. tracheiphila*-cucurbit host-insect vector pathosystem represents a valuable model study to understand the molecular, ecological, and evolutionary determinants that drive pathogen emergence due to agricultural modernization and other human-related activities.

## Methods

### Strains, culture media, growth conditions

Strains used in this study are listed in Supplementary Table 1. A spontaneous rifampicin resistant variant of *E. tracheiphila* strain BHKY (WT) (Rojas et al. 2013) and derived strains were used throughout the study. We used *Escherichia coli* strains TOP10 and Pir1 for cloning, preservation and propagation of gene constructions. For conjugation, *E. coli* strain S17-1λ was used as donor. *E. tracheiphila* was grown routinely in KB liquid media or agar at room temperature (25 ± 2°C). For induction of *hrpJ*, induction media was prepared as follows: 0.5 mM of K_2_SO_4_, 0.5 mM of CaCl_2_, 0.5 mM of Morpholine Ethane Sulfonic Acid (MES), 175 mM of mannitol, 5 mM of (NH_4_)_2_SO_4_, pH 7. *E. coli* strains were grown in LB liquid media or agar at 37°C. Antibiotics were used, when indicated, at the following concentrations: rifampicin, 50 μg/ml; ampicillin, 100 μg/ml; carbenicillin, 100 μg/ml; chloramphenicol, 5 μg/ml, kanamycin, 50 µg/ml.

### Scanning Electron Microscopy

Squash plants were inoculated with a culture of *E. tracheiphila* BHKY WT and maintained at greenhouse conditions. At 15 DPI, the stem from a plant showing wilting symptoms in the inoculated leaf was harvested 5 centimeters from the base of the leaf and cross-sectioned into 1 mm slices. Stem cross-sections were transferred to 4% paraformaldehyde in PBS buffer for 8 hours at room temperature to fix the samples. The samples were then washed three times with PBS buffer, followed by incubation for 10 minutes each in a series of ethanol solutions for dehydration (50, 70, 80, 90, 95% ethanol, and twice with 100% percent ethanol). The dehydrated samples were critical point dried, mounted on metal stubs with adhesive carbon tabs and sputter coated with a 2 nm layer of platinum/palladium 80:20. The samples were imaged immediately after sputter-coating at the Harvard Center for Nanoscale Systems using a Zeiss Supra 55VP field Emission SEM.

### Construction of deletion mutants

We generated deletion mutants by allelic replacement in operons *eps* and *hrp*T3SS (Figure 1, Supplementary Table S1). This was done by double homologous recombination with the *bla* gene, as described previously (Rocha et al. 2020). Specific primers sequences for the constructions are shown in Supplementary Table S3. Genetic constructions were assembled for deletion of *eps* and *hrp*T3SS operons, consisting of ≈0.8 kb fragments upstream and downstream the target locus flanking the *bla* cassette coding for Beta Lactamase. The *bla* gene was amplified using primers LS23 and LS24 from pDK46 (Datsenko and Wanner 2000). Flanking fragments were amplified from *E. tracheiphila* BHKY genomic DNA: for the Δ*eps* construction, the upstream fragment was amplified using primer pair JR75 and JR77, whereas for downstream fragment we used primer pair JR78 and JR80; for the Δ*hrp*T3SS construction, we used primer pair JR93 and JR95 for upstream region, and JR96 and JR98 for downstream region (Supplementary Table S3). After Gibson Assembly, reactions containing the constructions for Δ*eps* and Δ*hrp*T3SS were diluted 1:200 and re-amplified with primer pairs JR76-JR79 and JR94-JR97, respectively. Conjugation into *E. tracheiphila* BHKY WT and mutant selection was carried out as described previously (Rocha et al. 2020). Colonies of BHKYΔ*eps* and BHKYΔ*hrp*T3SS, (Supplementary Table 1) were grown in liquid media and stored at −80°C with 15% glycerol.

### Transcriptional fusions

GFP transcriptional-fusions were constructed for studying the expression of *amsG* and *hrpJ* genes from the *eps* and *hrp*T3SS loci, respectively (Fig. 1a), as well as *exlx* gene coding for Expansin (Rocha et al. 2020), using the plasmid pPROBE-GFP[AAV](Miller 2000). For studying gene expression *in planta*, this plasmid was modified by inserting in the *Sal*I site, a constitutively expressed *mcherry* gene, amplified from pMP7605 with primers JR138 and JR139, (Supplementary Table S3) generating plasmid pJR169 (Supplementary Table S1). Promoter region of the target genes were amplified using primers JR121 and JR59 for P*_amsG_* (585 pb), JR52 and JR53 for P*_hrpJ_* (281 pb) and JR126 and JR127 for P*_exlx_* (324 pb) (Supplementary Table S3). The promoters were inserted in the corresponding restriction sites of pJR169, and the resulting plasmids were transformed into chemically competent *Ec*-S17-1λ cells. These strains were used as donors for conjugation of *Et*-BHKY and conjugants were selected using KB plates with rifampicin and kanamycin. The resulting reporter strains display constitutive mCherry fluorescence as well as GFP fluorescence directed by the corresponding promoter.

### Virulence of *Erwinia tracheiphila* BHKY and derived strains in squash

Colonies of BHKY wildtype (WT) or mutant strains were picked for inoculating 3 ml of KB with the corresponding antibiotic (Supplementary Table S1) and grown for 48 h with shaking. From these cultures, 10 μl containing ≍10^7^ cells were used for inoculating 2-3 weeks old squash plants, in wounds induced in the petiole. Plants were kept in a greenhouse at room temperature with day/night cycles of 12/12 h. For co-inoculations, the corresponding stationary cultures were mixed 1:1 (v:v), and then 10 μl were used for inoculation. To assess disease development, we followed daily the apparition of wilt symptoms, and recorded the day of plant collapse or death, in days post-inoculation (DPI).

### Plant colonization experiments

For measuring bacterial growth *in planta*, samples of stems from plants inoculated with *E. tracheiphila* were sampled at each time point. Thin slices (<1 mm) of stem were obtained with a sterile blade and weighted. Then, 500 μl of sterile PBS were added and the stem slices were left in the buffer for 40 min in ice, vortexing every 10 min. Next, 100 μl of the suspension were used for 10-fold serial dilutions, which were plated in KB agar with rifampicin and the corresponding antibiotic (Supplementary Table S1). Colonies were counted after 5 days and CFU per fresh weight of plant was determined.

### Fluorescence microscopy

Two-week old squash plants were inoculated, as described above, using reporter strains *E. tracheiphila* WT (P*_amsG_’gfp*) or WT (P*_hrpJ_’gfp*) (Supplementary Table S1). These strains carry transcriptional fusions between *gfp* and promoters from the *eps* and *hrp*T3SS operons, respectively, as well as a constitutively expressed mCherry gene. At 15 DPI, plants were sampled for confocal microscopy. For this, the inoculated petiole was dissected manually into thin transversal sections of <1mm, which were briefly washed with sterile distilled water and dried. Next, samples were submerged into 0.5% low melting point agarose containing Calcofluor White (0.002 g/ml) (Mitra and Loqué 2014; Hughes and McCully 1975), and incubated at room temperature for 10 minutes. Agarose-covered samples were placed on coverslips and imaged using a Nikon Eclipse TE2000-U microscope equipped with a 60× Plan Apo oil objective, a Hamamatsu digital camera model ORCA-ER and using CFP/YFP (Chroma #52017), GFP (Chroma #41020) or Texas Red (Chroma #62002v2) filter sets.

### In vitro expression assays

For expression in vitro, colonies of reporter strains WT (P*_amsG’gfp_*), WT (P*_hrpJ’gfp_*) or a control carrying a promoterless *gfp* (Supplementary Table S1) were picked to inoculate 3 ml of KB with kanamycin and grown for 48 h to stationary phase. Then, cell pellets were obtained by centrifugation, washed with 1 volume of PBS and resuspended in 10X volume of KB or Induction Media (Wei et al. 1992). Replicates of 300 μl aliquots were placed in 96-well black microplates (Costar cat. 3916) and fluorescence was followed by exciting at 480 nm and detecting emission at 510 nm in a multimodal plate reader (SpectraMax M2, Molecular Devices) with controlled temperature.

### Flow cytometry

We analyzed *in planta* gene expression with a single cell approach. Plants were inoculated with reporter strains and cell suspensions were obtained for three biological replicates of each treatment at each time point, using the sampling method described above. Cell suspensions were analyzed by flow citometry using an LSR-II Analyser (BD-Biosciences, San Jose CA) for GPF and MCherry. The assay was normalized for recording 20,000 events of MCherry expressing cells, from which the percent of GFP expressing cells was obtained.

### Statistics

Statistical analyses were performed using the software PRISM 7 (GraphPad Software, La Jolla California USA, www.graphpad.com). Curves following symptoms in first leaf and plant death were compared using the built-in Log-rank (Mantel-Cox) test for survival analysis. When significant differences were found (*P* < 0.05), pairwise comparisons were tested using the same test (Machin et al. 2006).

## Acknowledgments

We thank all members of the Kolter lab for valuable discussion. We thank Chad Araneo for technical assistance at the Division of Immunology’s Flow Cytometry Core Facility. We also thank the Harvard Center for Nanoscale Systems, the facility where we conducted our electron microscopy studies. Dominique Schneider kindly donated the plasmid pDS132. This work was supported by Fundación Mexico en Harvard, and Conacyt grant 237414 to J.R., NSF postdoctoral fellowship DBI-1202736 to L.R.S. and NIH Grant GM58213 to R.K.

