## Supplementary Material for "Complementary roles of EPS, T3SS and Expansin for virulence of *Erwinia tracheiphila*, the causative agent of cucurbit wilt"

Supplementary Table S1. Strains and plasmids used in this study

| **Name** | **Description** | **Reference** |
|---|---|---|
| **Strains** | | |
| *Erwinia tracheiphila* BHKY (WT) | *Erwinia tracheiphila* Wt strain; $Rif^R$ | Rojas et al., 2013 |
| *E. tracheiphila* Δ*eps* | Derived from BHKY Wt; 17 Kb deletion in the *eps* operon; $Rif^R$, $Ap^R$ | This study |
| *E. tracheiphila* Δ*hrp*T3SS | Derived from BHKY; 15 kB deletion in the *hrp* Type III secretion system operon; $Rif^R$, $Ap^R$ | This study |
| *E. tracheiphila* Δ*exlx-gh5* | Derived from BHKY; deletion in the operon coding for Expansin-Endoglucanase; $Rif^R$, $Ap^R$. | Rocha et al., 2020 |
| *E. tracheiphila* Wt (pJR169) | BHKY Wt strain with the pJR169 plasmid, carrying a constitutiveluy expressed *mCherry* and a promoterless *gfp* | This study |
| *E. tracheiphila* Wt ($P_{amsG}$'*gfp*) | BHKY Wt reporter for EPS gene expression; carrying a constitutive *mCherry* and a $P_{amsG}$'*gfp* transcriptional fusion | This study |
| *E. tracheiphila* Wt ($P_{hrpJ}$'*gfp*) | BHKY Wt reporter for T3SS gene expression; carrying a constitutive *mCherry* and a $P_{hrpJ}$'*gfp* transcriptional fusion | This study |
| *E. tracheiphila* Wt ($P_{exlx}$'*gfp*) | BHKY Wt reporter for Expansin gene expression; carrying a constitutive *mCherry* and a $P_{exlx}$'*gfp* transcriptional fusion | This study |
| Top10 | *E. coli* strain used for general cloning | |
| PIR1 | *E. coli* strain for expression of R6K replication origin | |
| S17-1λ | *E coli* strain used as donor for conjugation | |

| **Plasmids** | | |
|---|---|---|
| pKD46 | Template for *bla* gene (ampicillin resistance cassette) | Datsenko and Wanner, 2000. |
| pJN105 | Low copy number plasmid | Newman and Fuqua, 1999. |
| pDS132 | Suicide plasmid for allelic replacement | Philippe et al., 2004 |
| pJN105cm | Derived from pJN105, modified with chloramphenicol resistance, used for genetic complementation | This study |
| pPROBE'GFP(AAV) | Plasmid for *gfp* transcriptional fusions | Miller et al., 2000 |
| pMP7605 | Template for PCR amplification of *mcherry* gene | Lagendijk et al., 2010 |
| pJR169 | Derived from pPROBE'GFP(AAV); modified with a constitutively expressed *mCherry* gene. | This study |

Supplementary Table S2. Statistical analyses of virulence assays with co-inoculations

| Comparison | Chi square | P value | Significant? (alpha 0.05) |
|---|---|---|---|
| **Symptoms on first leaf** | | | |
| All | 50.44 | <0.0001 | Yes |
| Wt vs Δ*eps* + Δ*hrp*T3SS | 25.21 | <0.0001 | Yes |
| Wt vs Δ*eps* + Δ*exlx-gh5* | 12.17 | 0.0005 | Yes |
| **Wt vs Δ*hrp*T3SS + Δ*exlx-gh5*** | **1.904** | **0.1637** | **No** |
| Wt vs Δ*exlx-gh5* | 23.06 | <0.0001 | Yes |
| Δ*exlx-gh5* vs Δ*eps* + Δ*exlx-gh5* | 0.7572 | 0.3842 | No |
| Δ*exlx-gh5* vs Δ*hrp*T3SS + Δexlx-gh5 | 15.35 | <0.0001 | Yes |
| **Death of plants** | | | |
| All groups | 32.57 | <0.0001 | Yes |
| Wt vs Δ*eps* + Δ*hrp*T3SS | 14.25 | 0.0002 | Yes |
| Wt vs Δ*eps* + Δ*exlx-gh5* | 8.737 | 0.0031 | Yes |
| Wt vs Δ*hrp*T3SS + Δ*exlx-gh5* | 5.283 | 0.0215 | Yes |
| Wt vs Δ*exlx-gh5* | 14.25 | 0.0002 | Yes |
| Δ*exlx-gh5* vs Δ*eps* + Δ*exlx-gh5* | 2.09 | 0.1483 | No |
| **Δ*exlx-gh5* vs Δ*hrp*T3SS + Δ*exlx-gh5*** | **4.609** | **0.0318** | **Yes** |

Supplementary Table 3. Oligonucleotides used in this study.

| Code | Sequence | Restriction site or modification |
|---|---|---|
| LS23 | acttttcggggaaatgtgc | |
| LS24 | acgttaagggattttggtca | |
| ***eps* operon deletion** | | |
| JR75 | attgctcgaggatttgatc | |
| JR76 | gaggagctcgcaaatgaccgaaatactcc | *Sac*I site |
| JR77 | gcacatttccccgaaaagtcttgcgataggtatagtgg | LS23 tag |
| JR78 | tgaccaaaatcccttaacgtgttgcgtatcaaatttctcc | LS24 tag |
| JR79 | gaggagctcccaaaacctgttgatgacc | *Sac*I site |
| JR80 | cctgtggtcttattgtctgg | |
| ***hrpT3SS* deletion** | | |
| JR93 | cctctacttccgtcttaacacc | |
| JR94 | gaggagctcctgtttcgtacattcgtgg | *Sac*I site |
| JR95 | gcacatttccccgaaaagtcagaactgaatggatttagagg | LS23 tag |
| JR96 | tgaccaaaatcccttaacgtaggagtgacatgatgacacc | LS24 tag |
| JR97 | gaggagctcgagaatatcgatacccacacg | *Sac*I site |
| JR98 | ccagtgtctgatgattgttgc | |
| **Transcriptional fusions** | | |
| JR138 | gtcgtcgacgagctgttgacaattaatcatcggctcg | *SalI* |
| JR139 | gtcgtcgacgttttttcgttttcttacttgtacag | *SalI* |
| JR121 | ggaggatccgcaaatgaccgaaatactcc | *BamH*I |
| JR59 | gaagaattcaatagcctgtgaataattacttcc | *EcoR*I |
| JR52 | ggaggatccactgttctgtggactgcaag | *BamH*I |
| JR53 | gaagaattccaatgtgtggcgtgattag | *EcoR*I |
| JR126 | ggaggatccccagcataagctcaactcag | *BamH*I |
| JR127 | gaagaattcgataaaggtgactgagaaagtcag | *EcoR*I |

Supplementary Figure S1

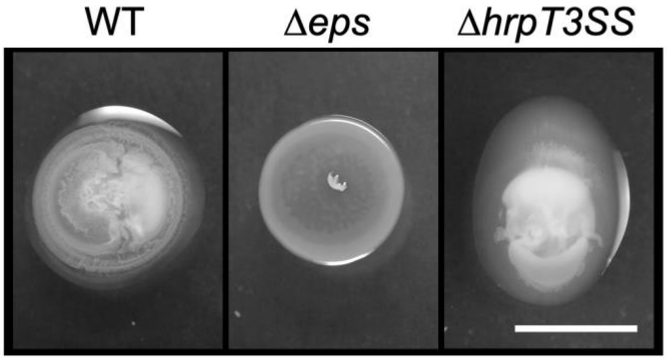


Supplementary Figure S1. Effect of gene deletions on the mucoid colony phenotype of *E. tracheiphila* in vitro. Five µl of each strain were spot-inoculated on the surface of KB agar and grown for 5 days at 25 ˚C. a) Effect of operon deletions; b) effect of frameshift deletion in *amsG* gene and genetic complementation. Scale bar, 5 mm.

Supplementary Figure S2

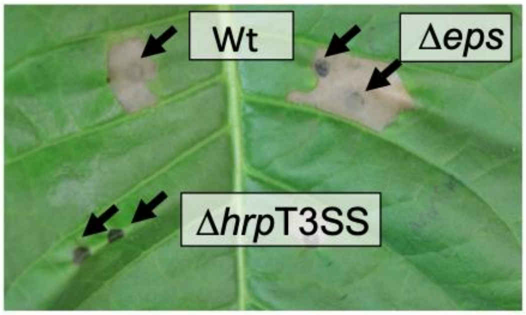


Supplementary Figure S2. Effect of gene deletion on the hyper sensitivity response (HR) caused by *E. tracheiphila* on tobacco. Strains were grown overnight in KB and 300 μl were used to infiltrate the leaf through the lower side with a needleless syringe. Leaves were imaged 2 days after inoculation. Arrows indicate the sites of infiltration.

Supplementary Figure S3

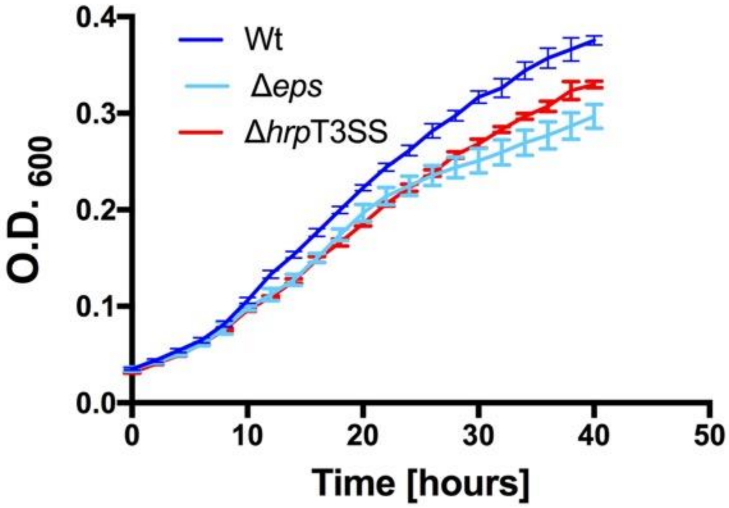


Supplementary Figure S3. In vitro growth of *E. tracheiphila* WT, Δ*eps* and Δ*hrp*T3SS strains in KB media. Overnight cultures were washed and cells were diluted to OD600 of 0.03 in fresh KB. Diluted cultures were pipetted in four replicates into 96 well plates and absorbance was measured every 2 h for 40 h at 25˚C.

Supplementary Figure S4

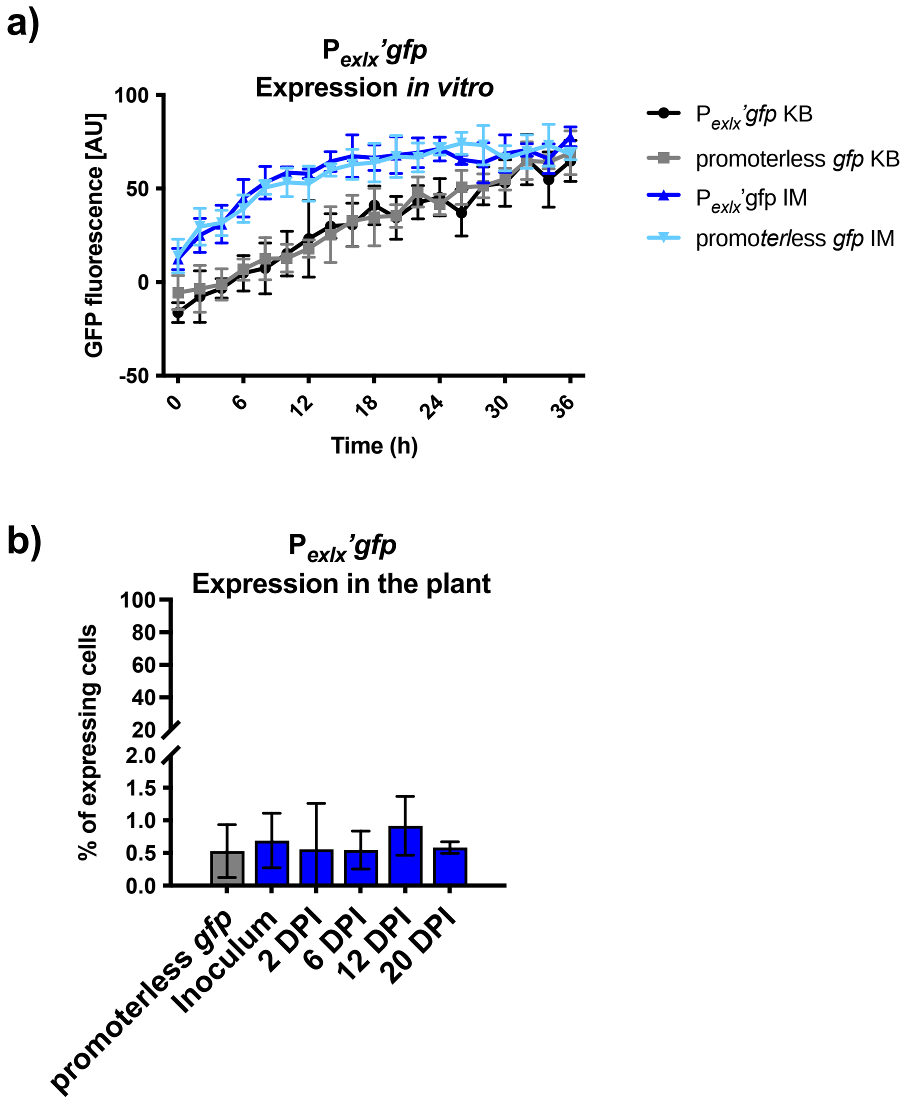


Figure S4. Expression of expansin was not detected form the reporter strain *E. tracheiphila* WT $P_{exlx}$'*gfp* a) in vitro or b) during squash infection. a) Reporter strain carrying the transcriptional fusion $P_{exlx}$'*gfp* and the control strain with promoterless *gfp* were inoculated at OD600 of 0.03 in a microplate in KB rich media or Induction Media (IM), and GFP fluorescence was followed for 36 h. b) Squash plants were inoculated with the reporter strain *E. trachieiphila* ($P_{exlx}$'*gfp*) and samples were collected for flow cytometry analysis throughout the infection for quantification of GFP-expressing cells.

Supplementary Figure S5

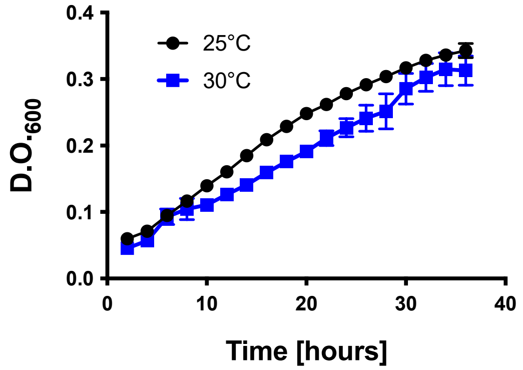


Figure S4. In vitro growth is not severely affected by increased temperature. *E. tracheiphila* WT was grown in KB rich media, and OD600 was followed for 36 h. Average and standard deviation of four replicates is shown.
